# Proteomic profiling of the local and systemic immune response to pediatric respiratory viral infections

**DOI:** 10.1101/2024.10.08.617294

**Authors:** Emily Lydon, Christina M. Osborne, Brandie D. Wagner, Lilliam Ambroggio, J. Kirk Harris, Ron Reeder, Todd C. Carpenter, Aline B. Maddux, Matthew K. Leroue, Nadir Yehya, Joseph L. DeRisi, Mark W. Hall, Athena F. Zuppa, Joseph Carcillo, Kathleen Meert, Anil Sapru, Murray M. Pollack, Patrick McQuillen, Daniel A. Notterman, Charles R. Langelier, Peter M. Mourani, Eunice Kennedy Shriver National Institute of Child Health and Human Development Collaborative Pediatric Critical Care Research Network (CPCCRN)

## Abstract

Viral lower respiratory tract infection (vLRTI) is a leading cause of hospitalization and death in children worldwide. Despite this, no studies have employed proteomics to characterize host immune responses to severe pediatric vLRTI in both the lower airway and systemic circulation. To address this gap, gain insights into vLRTI pathophysiology, and test a novel diagnostic approach, we assayed 1,305 proteins in tracheal aspirate (TA) and plasma from 62 critically ill children using SomaScan. We performed differential expression (DE) and pathway analyses comparing vLRTI (n=40) to controls with non-infectious acute respiratory failure (n=22), developed a diagnostic classifier using LASSO regression, and analyzed matched TA and plasma samples. We further investigated the impact of viral load and bacterial coinfection on the proteome. The TA signature of vLRTI was characterized by 200 DE proteins (P_adj_<0.05) with upregulation of interferons and T cell responses and downregulation of inflammation-modulating proteins including FABP and MIP-5. A nine-protein TA classifier achieved an AUC of 0.96 (95% CI 0.90-1.00) for identifying vLRTI. In plasma, the host response to vLRTI was more muted with 56 DE proteins. Correlation between TA and plasma was limited, although ISG15 was elevated in both compartments. In bacterial coinfection, we observed increases in the TNF-stimulated protein TSG-6, as well as CRP, and interferon-related proteins. Viral load correlated positively with interferon signaling and negatively with neutrophil-activation pathways. Taken together, our study provides fresh insight into the lower airway and systemic proteome of severe pediatric vLRTI, and identifies novel protein biomarkers with diagnostic potential.

**IMPORTANCE:** We describe the first proteomic profiling of the lower airway and blood in critically ill children with severe viral lower respiratory tract infection (vLRTI). From tracheal aspirate (TA), we defined a proteomic signature of vLRTI characterized by increased expression of interferon signaling proteins and decreased expression of proteins involved in immune modulation including FABP and MIP-5. Using machine learning, we developed a parsimonious diagnostic classifier that distinguished vLRTI from non-infectious respiratory failure with high accuracy. Comparative analysis of paired TA and plasma specimens demonstrated limited concordance, although the interferon-stimulated protein ISG15 was significantly upregulated with vLRTI in both compartments. We further identified TSG-6 and CRP as airway biomarkers of bacterial-viral coinfection, and viral load analyses demonstrated positive correlation with interferon-related protein expression and negative correlation with the expression of neutrophil activation proteins. Taken together, our study provides new insight into the lower airway and systemic proteome of severe pediatric vLRTI.

## INTRODUCTION

Respiratory viral infections are the most common cause of pediatric illness worldwide.^1^ While often mild and self-limited, a substantial number of children progress to severe viral lower respiratory tract infection (vLRTI) requiring hospital admission and mechanical ventilation (MV), often further complicated by acute respiratory distress syndrome (ARDS) and/or bacterial coinfections. In a global epidemiological study of children under five, severe LRTI was the leading cause of mortality outside of the neonatal period, contributing to an estimated 760,000 deaths.^2^

The marked heterogeneity in vLRTI clinical outcomes, driven in large part by differential host responses, remains poorly understood.^3^ Deeply profiling the host immune response to vLRTI can offer insights into pathophysiology and also enable novel diagnostic test development.^4,5^ Prior work evaluating the host response in LRTI using systems biology approaches has mainly focused on adult populations, and the few pediatric LRTI studies predominantly utilized transcriptomic or metabolomic approaches.^6–10^ Proteomics, or the large-scale study of the protein composition within a biologic sample, has the potential to complement studies of the transcriptome, as protein expression is influenced by post-transcriptional regulation and may be a more direct reflection of cellular and immunologic processes.^11^ The limited number of proteomic pediatric LRTI studies published to date have profiled plasma or urine samples,^12–14^ which provide useful insights into the systemic response to LRTI and offer candidate diagnostic biomarkers, but may not reflect biological processes at the site of active infection. The local host proteomic response to severe viral infection in the lower respiratory tract remains poorly understood in children, as does the compartmentalization of proteomic responses in the blood versus airway.

To address these questions, we perform high-dimensional proteomic profiling of paired tracheal aspirate (TA) and plasma samples in a prospective multicenter cohort of critically ill children with acute respiratory failure, specifically comparing vLRTI to non-infectious etiologies. We hypothesized that there would be a distinct proteomic signature of vLRTI, more pronounced in the airway than blood, and that exploring proteomic correlations with bacterial-viral coinfection and viral load would yield valuable biological insights.

## METHODS

### Description of cohort

Children in this study represent a subset of those enrolled in a previously described prospective cohort of 454 mechanically ventilated children admitted to eight pediatric intensive care units in the National Institute of Child Health and Human Development’s (NICHD) Collaborative Pediatric Critical Care Research Network (CPCCRN) from February 2015 to December 2017.^8,9^ See supplementary material for enrollment criteria. IRB approval was granted for TA sample collection prior to consent, as endotracheal suctioning is standard-of-care. Specimens of children for whom consent was not obtained were destroyed. The study was approved by University of Utah IRB #00088656.

### Sample collection and processing

TA specimens collected within 24 hours of intubation were processed for proteomic analysis, with centrifugation at 4°C at 15,000xg for five minutes and freezing of supernatant at -80°C in a microvial within 30 minutes. Some patients did not have TA samples available for proteomic analysis due to inadequate processing. Plasma samples collected within 24 hours of MV were frozen at -80°C. Some patients did not have plasma collected because consent was not obtained within the timeframe.

### Adjudication of LRTI status

Adjudication was carried out retrospectively by study-site physicians who reviewed all clinical, laboratory, and imaging data following hospital discharge, with specific criteria detailed in the supplementary material. Standard of care microbiological testing, including multiplex respiratory pathogen polymerase chain reaction (PCR) and semiquantitative bacterial respiratory cultures, were considered in the adjudication process. In addition, microbes detected by TA metagenomic next-generation sequencing (mNGS), as previously described,^8^ were considered for pathogen identification. Patients were assigned a diagnosis of “vLRTI” if clinicians made a diagnosis of LRTI and the patient had a respiratory virus detected by PCR and/or mNGS. Within the vLRTI group, subjects were subcategorized as viral infection alone or bacterial coinfection, based on whether a bacterial respiratory pathogen was detected by bacterial culture, PCR, and/or mNGS. Patients alternatively were assigned a diagnosis of “No LRTI” if clinicians identified a clear, non-infectious cause of respiratory failure without clinical or microbiologic evidence of bacterial or viral LRTI.

### Subject selection for proteomic analysis

Subjects that were clinically adjudicated as vLRTI and No LRTI were selected for proteomics analysis, in an approximately 2:1 ratio. This subset of subjects represented a convenience sample of the larger cohort, with the goal of maximizing the number of subjects with both TA and plasma samples to allow comparative proteomic analysis, although not all subjects had all both samples available.

### Proteomic analysis

The SomaScan^®^ 1.3k assay (SomaLogic) was utilized to quantify the protein expression in plasma and TA samples. The assay, described and validated elsewhere,^15–17^ utilizes 1,305 single-stranded DNA aptamers that bind specific proteins, which are quantified on a customized Agilent hybridization assay. Aptamer measurement is therefore a surrogate of protein expression. The assay outputs fluorescence units that are relative but quantitatively proportional to the protein concentration in the sample.

### Statistical analysis

Relative fluorescence units (RFUs) for each of the 1,305 protein aptamers were log-transformed for analysis. Differential expression was calculated between groups for each aptamer using limma, a R package that facilitates simultaneous comparisons between numerous targets.^18^ Age-adjusted and age-unadjusted differential protein analyses were performed. Biological pathways were interrogated against the Reactome database with the R package WebGestaltR using a functional class scoring approach.^19,20^ Specifically, the input list included the full set of 1,305 proteins and the corresponding log2-fold change between the conditions of interest, ranked by T-statistic. P-values for protein and pathway analyses were adjusted for multiple comparisons using the Benjamini-Hochberg procedure; adjusted p-value (p_adj_)<0.05 was considered statistically significant.

A parsimonious proteomic classifier was generated using LASSO logistic regression on TA samples with the cv.glmnet function in R, setting family = “binomial” and leaving other parameters as default.^21^ LASSO was used for both feature selection and classification. The model was generated using five-fold cross-validation, where a model was trained on ∼80% of samples and tested on ∼20% of samples to generate vLRTI probabilities for each of the subjects in the cohort. To keep the fold composition comparable, we required at least 3 No LRTI subjects in each fold. AUC was calculated using the pROC package and confidence intervals were generated with bootstrapping.^19^

Correlation for each protein between TA and plasma samples were calculated using Pearson correlation, for all paired samples in bulk and then subdivided by group (vLRTI vs No LRTI). Correlation coefficients were considered strong if the absolute value was >0.5, moderate if 0.3-0.5, and weak if 0-0.3. Correlation between specific proteins and viral load were calculated similarly. Viral load was extrapolated from mNGS reads-per-million, and if multiple viruses were detected, the viral loads were summed.

## RESULTS

### Cohort characteristics and microbiology

From the prospective multi-center cohort (n=454), samples from 62 subjects underwent proteomic analysis, including 40 with vLRTI and 22 with No LRTI (**Figure 1A**). Those with vLRTI were further subdivided into viral infection alone (n=16) or viral-bacterial coinfection (n=24). The demographic characteristics did not differ between the vLRTI and No LRTI groups, with the exception of age, which was higher in No LRTI than vLRTI (median 10.2 years [IQR 1.1-14.9] vs 0.9 [0.3-1.6]) (**Table 1**). Diagnoses in the No LRTI group included trauma, neurologic conditions, ingestion, and anatomic airway abnormalities, with many having abnormal chest radiographs and meeting ARDS criteria. Within the vLRTI group, respiratory syncytial virus (RSV) was the most common pathogen, and 15 subjects had more than one virus (**Figure 1B**). *Haemophilus influenzae, Moraxella catarrhalis*, and *Streptococcus pneumoniae* were most common coinfecting bacterial pathogens.

**Table 1:** Demographic and clinical characteristics of the vLRTI and No LRTI cohorts.

|  | **vLRTI (n=40)** | **No LRTI (n=22)** | **P value\*** |
| --- | --- | --- | --- |
| **Female, n (%)** | 18 (45.0%) | 11 (50.0%) | 0.79 |
| **Age in years, median (IQR)** | 0.9 (0.3 - 1.6) | 10.2 (1.1-14.9) | <0.01 |
| **Race** |  |  | 0.71 |
| **White, n (%)** | 26 (65.0%) | 14 (63.6%) |  |
| **Black/African American, n (%)** | 6 (15.0%) | 6 (27.3%) |  |
| **Asian, n (%)** | 1 (2.5%) | 1 (4.5%) |  |
| **Native Hawaiian/Pacific Islander, n (%)** | 1 (2.5%) | 0 (0.0%) |  |
| **American Indian/Alaska Native, n (%)** | 1 (2.5%) | 0 (0.0%) |  |
| **Unknown/Not Reported, n (%)** | 5 (12.5%) | 1 (4.5%) |  |
| **Hispanic/Latino ethnicity, n (%)** | 8 (20.0%) | 1 (4.5%) | 0.14 |
| **Comorbidity, n (%)** | 14 (35.0%) | 11 (50.0%) | 0.29 |
| **Immunocompromise, n (%)** | 1 (2.5%) | 0 (0.0%) | 0.99 |
| **Admission category** |  |  | <0.01 |
| **Medical, n (%)** | 40 (100%) | 14 (63.6%) |  |
| **Surgical, n (%)** | 0 (0.0%) | 5 (22.7%) |  |
| **Trauma, n (%)** | 0 (0.0%) | 3 (13.6%) |  |
| **Infiltrates on initial CXR, n (%)** | 36 (90.0%) | 12 (54.5%) | <0.01 |
| **ARDS, n (%)** | 18 (45.0%) | 4 (18.0%) | 0.05 |
| **Received antibiotics, n (%)** | 14 (35.0%) | 8 (36.4%) | 0.99 |
| **Ventilator days, median (IQR)** | 6 (5.0-9.0) | 6 (5.0-8.0) | 0.85 |
| **ICU length of stay in days, median (IQR)** | 10 (8.0-16.5) | 9 (6.3-13.3) | 0.25 |
| **Hospital length of stay in day, median (IQR)** | 14 (10.5-19.5) | 16 (9.5-38.8) | 0.34 |
| **Mortality, n (%)** | 1 (2.5%) | 3 (13.6%) | 0.12 |

\*Wilcoxon rank sum test used for all continuous variables. Fisher's exact test used for all categorical variables. IQR, interquartile range; ARDS, acute respiratory distress syndrome; ICU, intensive care unit.

**Figure 1:**
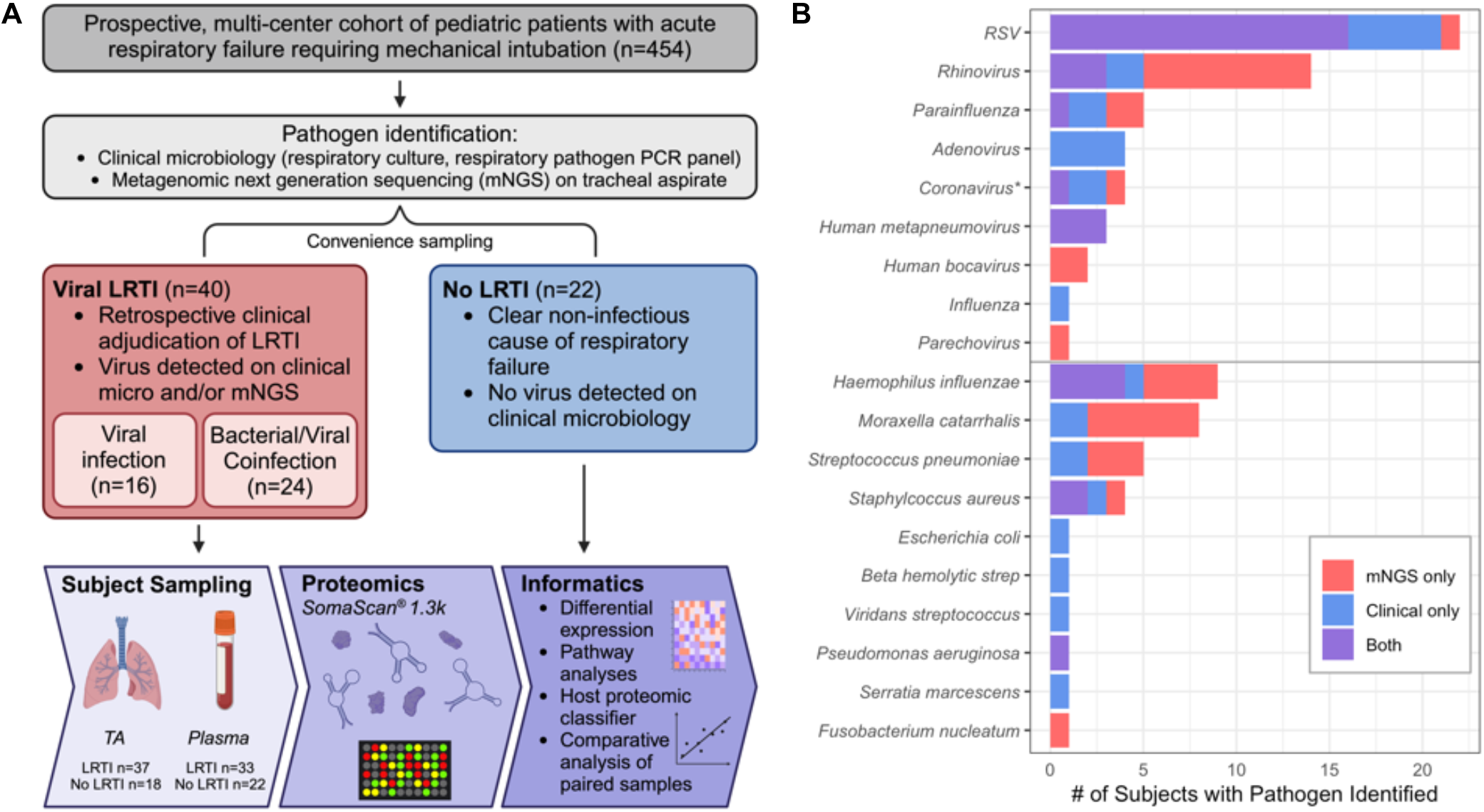
Study overview. A) From a prospectively enrolled multicenter cohort of pediatric patients presenting with acute respiratory failure requiring intubation, 62 were selected for proteomic analysis. The vLRTI group included a subset of subjects who had bacterial coinfection. Plasma and tracheal aspirate (TA) samples collected on enrollment underwent proteomic profiling on the SomaScan^®^ platform. Some subjects did not have all samples available for analysis, so the numbers available for each sample type are shown. Informatics approaches included evaluation of differentially expressed proteins and pathways, development of a host proteomic classifier, and comparative analyses of paired samples. B) Microbiology of the vLRTI group. Bar plot color indicates whether the microbe was detected on clinical microbiology, tracheal aspirate metagenomic next-generation sequencing (mNGS), or both. Many subjects had co-detection of multiple pathogens; thus, the total number of pathogens exceeds the number of patients in the vLRTI cohort. *Coronavirus includes only non-SARS-CoV-2 coronaviruses.

### Defining a lower respiratory tract proteomic signature of vLRTI

We first compared protein expression in TA samples between the vLRTI (n=37) and No LRTI (n=18) groups. Two hundred proteins (15.3% of all proteins assayed) were differentially expressed at p_adj_<0.05 (**Figure 2A, 2B**). Among the 80 proteins upregulated in vLRTI were interferon-stimulated ubiquitin-like protein ISG15 and oligoadenylate synthase protein OAS1, which are central to type I interferon signaling, and Granzyme B and Granulysin, proteins present in granules of cytotoxic T cells and natural killer (NK) cells. Among the 120 proteins downregulated in vLRTI were fatty acid-binding protein FABP, macrophage inhibitory protein MIP-5, and neutrophil-activating protein NAP-2. Pathway analysis confirmed interferon signaling as the primary pathway upregulated in the vLRTI group, although only the “influenza infection” pathway achieved p_adj_<0.05 (**Figure 2C**).

**Figure 2:**
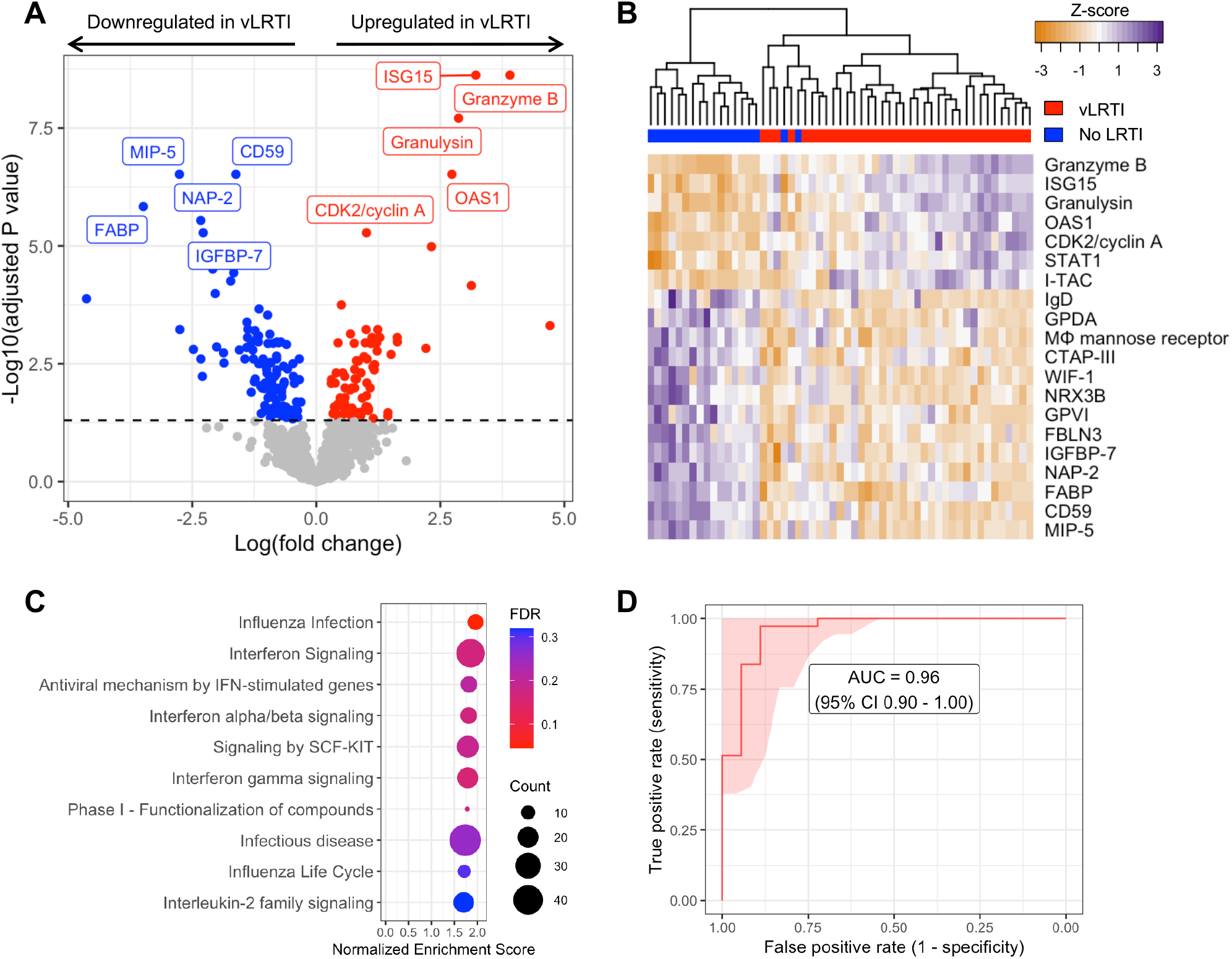
Comparison of host protein expression between vLRTI and No LRTI cohorts in tracheal aspirate. A) Volcano plot of the differentially expressed proteins, with proteins significantly upregulated in vLRTI in red, and proteins significantly downregulated in vLRTI in blue. The top ten proteins based on P_adj_ are labeled. B) Heat map showing differential expression of the top 20 proteins based on P_adj_ (rows) across all patients (columns). Dendrogram clustering (top) highlights the proteomic differences between the two groups. C) Pathway analysis showing the top ten pathways (all upregulated) ordered by Normalized Enrichment Score. Dot color indicates the false discovery rate (FDR) P_adj_, and size indicates the number of proteins included in the pathway. D) Receiver operator characteristic (ROC) curve of the proteomic classifier to distinguish vLRTI from No LRTI.

Having identified a strong host proteomic signature of vLRTI, we hypothesized that a parsimonious number of TA proteins could accurately differentiate vLRTI from No LRTI subjects. Utilizing LASSO logistic regression and employing five-fold cross-validation, we built parsimonious proteomic classifiers (ranging in size from 9 – 15 proteins) that accurately distinguished vLRTI and No LRTI with an area under the receiver operator curve (AUC) of 0.96 (95% CI 0.90-1.00) (**Figure 2D, Table S1**). The proteins with consistently positive coefficients (i.e. increasing vLRTI probability) were Granulysin, Granzyme B, and ISG-15 as well as cyclin-dependent kinase protein CDK2 and kinesin-like protein KIF23. The proteins with consistently negative coefficients (i.e. decreasing probability of vLRTI) were FABP and NAP-2.

Since age was statistically different between the two groups, we added age as a continuous covariate in our differential expression model (**Figure S1**). The results overall were similar, with 176 differentially expressed proteins (58 upregulated and 118 downregulated with vLRTI). There was considerable overlap (80%) in the top 10 most differentially expressed proteins between the two models.

### Comparison of plasma proteomics between vLRTI and No LRTI groups

We next compared plasma protein concentrations between vLRTI (n=33) and No LRTI (n=22) groups. The age-unadjusted differential expression analysis yielded 56 statistically significant proteins (4.3% of all proteins assayed), 45 upregulated in vLRTI and 11 downregulated in vLRTI (**Figure 3A**). However, adjusting for age, only one protein, ISG15, remained significant (**Figure S2**). ISG15, a type 1 interferon-stimulated protein, showed promise in distinguishing vLRTI and No LRTI groups in both plasma and TA (p_adj_ for both <0.0001), and ISG15 expression was strongly correlated between paired TA and plasma samples (correlation coefficient 0.79, p<0.0001) (**Figure 3B, 3C**). ISG-15 alone implemented as a diagnostic test exhibited strong performance with AUCs of 0.95 (95% CI 0.89-1.00) and 0.91 (95% CI 0.83-0.99) in TA and plasma, respectively (**Figure S3**).

**Figure 3:**
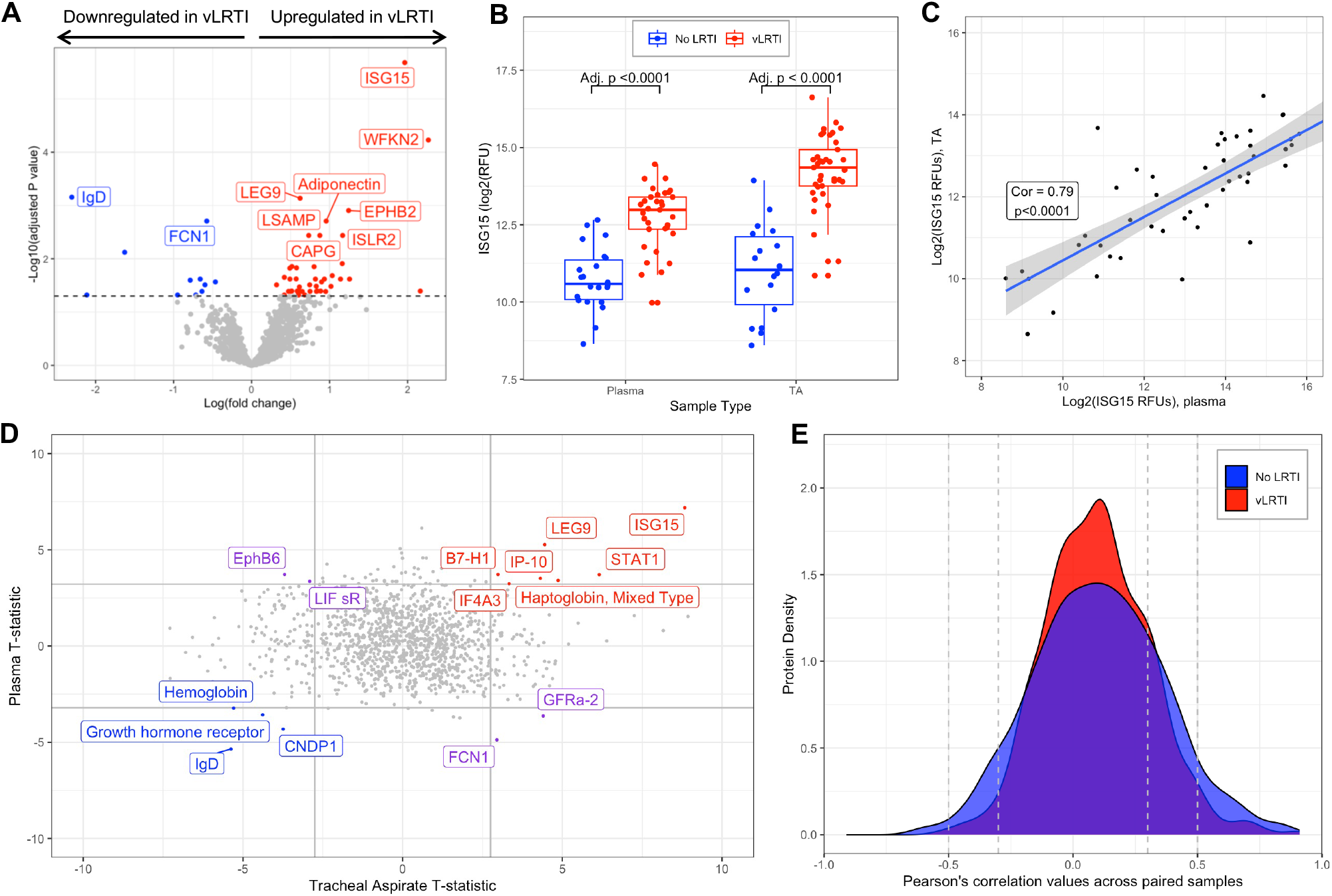
Host protein expression in plasma and comparative proteomic analysis between plasma and tracheal aspirate samples. A) Volcano plot of the differentially expressed plasma proteins in vLRTI versus No LRTI (age unadjusted). The top ten proteins based based on P_adj_ are labeled. B) Ubiquitin-like ISG-15 protein expression in plasma (left) and tracheal aspirate (TA) (right) in vLRTI (red) and No LRTI (blue) groups. C) Log-log plot showing correlation of ISG-15 values between paired plasma (x-axis) and TA (y-axis) samples. D) T statistics for each protein calculated with limma for vLRTI versus No LRTI comparisons in plasma and TA were plotted against one another. Proteins highlighted in red were significantly upregulated across both body compartments, in blue were downregulated in both, and in purple deviated in opposite directions. E) Density plot showing correlation coefficients for each protein in TA versus plasma, with stratification based on group (vLRTI in red vs No LRTI in blue).

### Comparative analysis of plasma and respiratory tract proteomics

Comparing the differentially expressed proteins between vLRTI and No LRTI groups in TA and plasma (using the age-unadjusted analyses), only 15 proteins were differentially expressed in both compartments, with seven proteins upregulated in vLRTI in both, four proteins downregulated in vLRTI in both, and four proteins with opposite directionality (**Figure 3D**). We further investigated protein correlation utilizing our paired samples (n=48 total paired TA and plasma samples from the same subject, including n=30 paired vLRTI samples and n=18 paired No LRTI samples). Correlation in expression between the lower airway and systemic circulation was weak for the majority of proteins (Pearson correlation coefficient -0.3 to +0.3) (**Figure 3E**), though there were exceptions, namely ISG-15.

### Lower respiratory tract proteomic differences in bacterial-viral coinfection

Within the vLRTI group, subjects were categorized as either viral infection (n=16) or bacterial-viral coinfection (n=24) based on clinical microbiology and respiratory mNGS. Differential protein expression in TA between these two groups did not yield any statistically significant proteins at p_adj_<0.05, but we did note an absolute increase in the expression of TSG-6, a tumor-necrosis factor-stimulated protein (p_adj_=0.07) and C-reactive protein (CRP) (p_adj_=0.10) in coinfection (**Figure 4A**). Pathway analysis showed heightened interferon signaling in coinfection compared to viral infection alone. Pathways associated with cell turnover and division were preferentially upregulated in viral infection compared to coinfection (**Figure 4B**).

**Figure 4:**
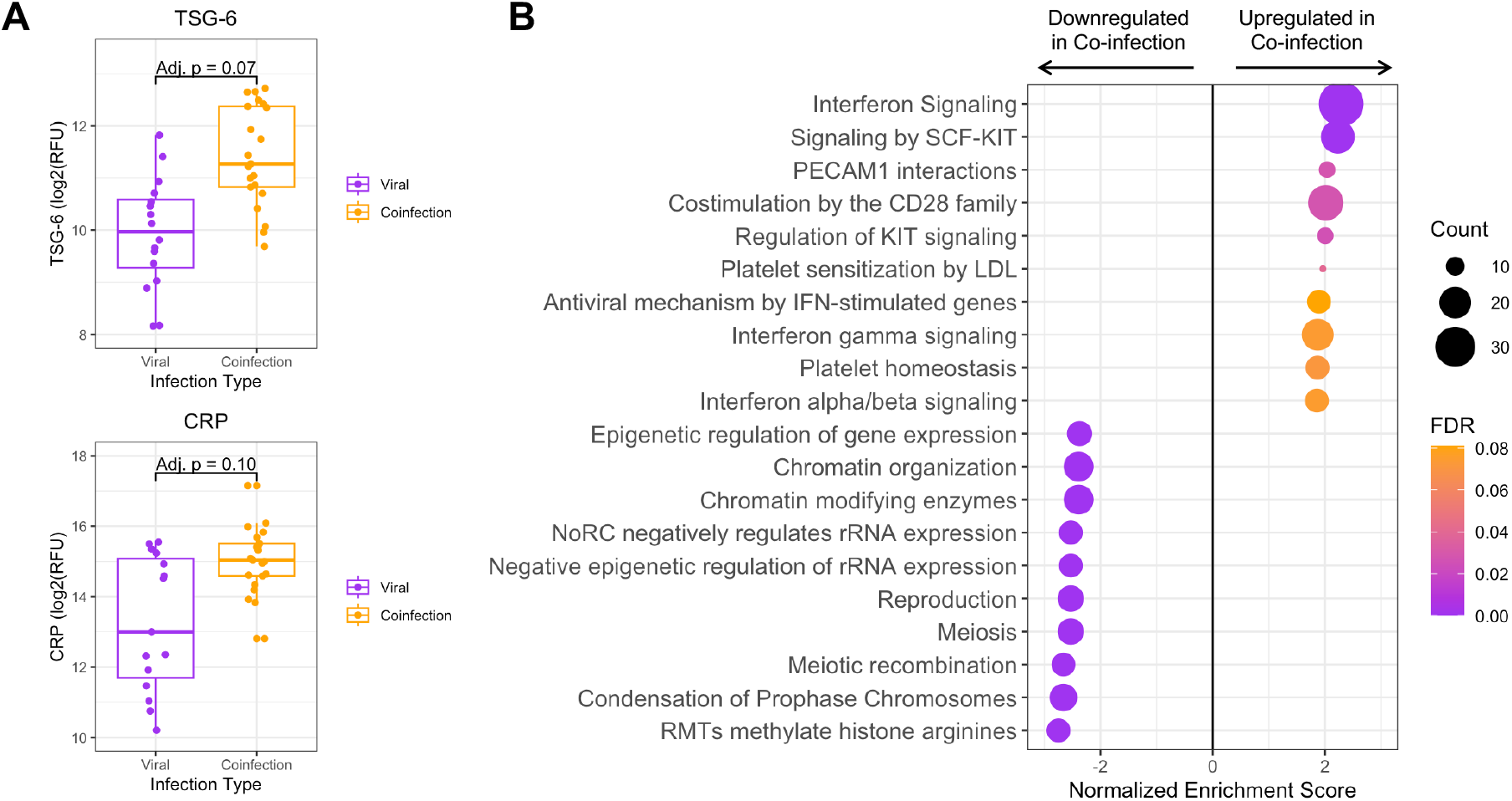
Tracheal aspirate protein and pathway expression in bacterial-viral coinfection. A) Box plots of the two most differentially expressed proteins, tumor necrosis factor stimulated gene-6 (TSG-6) and C-reactive protein (CRP), between viral infection and coinfection subgroups. B) Pathway analysis showing the top twenty pathways up- or down-regulated in bacterial-viral coinfection compared to viral infection alone. Dot color indicates the false discovery rate (FDR) P_adj_, and size indicates the number of proteins included in the pathway.

### Lower respiratory tract protein correlations with viral load

For the vLRTI subjects that tested positive for a virus by mNGS, the expression of TA proteins was correlated with viral load, measured as reads-per-million (**Figure 5**). Interferon-related proteins such as interferon-lambda 1 and ISG-15 were positively correlated with viral load, as well as monocyte chemotactic protein MCP-2. Conversely, platelet receptor GI-24, TFG-β superfamily protein Activin AB, and neutrophil-activating glycoprotein CD177 were inversely correlated with viral load.

**Figure 5:**
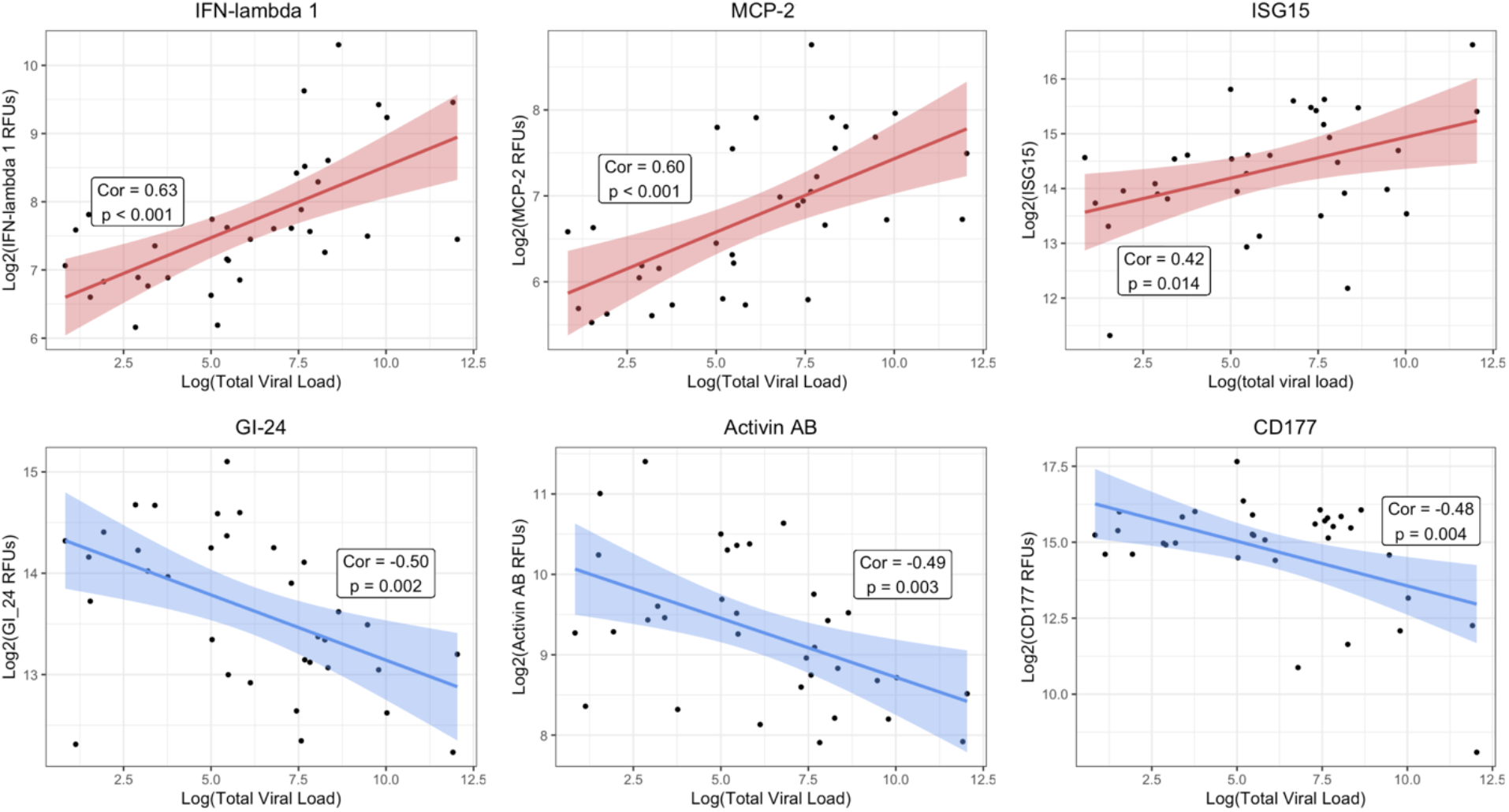
Correlation of lower respiratory protein expression and viral load. For each vLRTI subject with a virus detected on mNGS, correlation coefficients were calculated between viral load and relative concentration of each protein in TA. The top three highest positive correlations and the top three negative correlations are shown: interferon lambda-1 (IFN-lambda 1), monocyte chemotactic protein-2 (MCP-2); ubiquitin-like interferon stimulated gene-15 (ISG-15), platelet receptor GI-24, activin AB, and CD177.

## DISCUSSION

In this study, we identified the proteomic signature of severe pediatric vLRTI in both the lower respiratory tract and systemic circulation, leveraging results to understand compartment-specific host responses, host-viral dynamics, and viral-bacterial coinfection, as well as identify specific proteins with diagnostic potential. This work represents the first simultaneous proteomic profiling of both TA and plasma samples from children with severe vLRTI.

As hypothesized, the proteomic response to vLRTI was most robust at the local site of infection, with approximately 15% of assayed proteins differentially expressed in TA. This lower airway proteomic vLRTI signature was dominated by interferon-related proteins, which are well-known innate mediators of host defense and immunologic injury in viral infection.^22^ In addition, this signature was enriched in proteins contained in cytotoxic lymphocytes that in turn secrete interferons.^23^ Notable downregulated proteins were macrophage inhibitory protein-5 (MIP-5) and fatty acid binding protein (FABP), which has a diverse array of functions including macrophage regulation, suggesting that macrophage dynamics play an important role in response to vLRTI.^24,25^ Interestingly, in the subanalysis of bacterial-viral coinfection, the expression of interferon-related proteins was even greater than in viral infection alone. Prior work, mostly in influenza infections, has suggested that type 1 interferons can suppress key neutrophil and macrophage defenses, increasing susceptibility to bacterial coinfection, which may explain this finding.^26,27^

By integrating viral load measurements, we identified host TA proteins exhibiting proportional changes in expression based on viral load. Interferon-related proteins, including ISG-15 and interferon-lambda 1, and MCP-2, a chemokine induced by interferon signaling, exhibited the strongest induction in expression with viral load, underscoring the central role of interferons in innate antiviral defense. In contrast, the levels of CD177 (a glycoprotein involved in neutrophil activation),^28^ Activin AB (a TGF-β family protein implicated in ARDS inflammatory remodeling),^29^ and GI-24 (a platelet aggregation receptor)^30^ all decreased in response to higher viral loads. As previously noted, impaired neutrophil responses have been implicated in the pathophysiology of post-viral bacterial pneumonia,^26,27^ and our results suggest that this may occur in a viral-load dependent manner. Complementing these findings, a longitudinal transcriptomic study in adults hospitalized with severe influenza infection demonstrated initial upregulation of interferon pathways followed by inflammatory neutrophil activation and cell-stress patterns,^31^ and a study of severe pediatric influenza infection found that early upregulation of genes associated with neutrophil degranulation were associated with multi-organ dysfunction and mortality.^32^

While we observed a robust protein signature of vLRTI in the lower airways, the findings in the peripheral blood were more subtle, and correlation between plasma and TA proteins was generally weak. Furthermore, we observed a greater impact of age on the blood proteomic signature of vLRTI, potentially because the signal in the peripheral blood was weaker and thus more susceptible to confounding. Understanding the systemic response to a local infection is certainly useful and practical, as peripheral blood samples and urine samples are less invasive to collect than lower respiratory samples and would allow for application in a broader population of children who do not require MV. However, to obtain the most informative and potent proteomic signal of infection, our findings suggest that sampling the site of infection has the highest yield. Supporting this intuitive finding is a comparative adult proteomic study assaying both serum and bronchoalveolar lavage in interstitial lung diseases which similarly found a much higher number of differentially expressed proteins in the lower respiratory tract compared to the blood.^33^

In addition to insights into the pathophysiology of vLRTI, our study also highlights the utility of proteomic approaches in diagnostic biomarker discovery. Standard-of-care multiplexed PCR assays only evaluate a limited subset of respiratory viruses^34^ and cannot detect novel emerging viruses or differentiate asymptomatic carriage from true infection.^35^ Host response-based assays agnostic to viral species could be invaluable for pandemic preparedness and infection prevention in congregate settings. When employed as a diagnostic test to distinguish vLRTI from non-infectious respiratory failure, our nine-protein TA classifier achieved excellent performance with an AUC of 0.96. The single protein ISG15 also showed potential for use as a diagnostic biomarker in both TA and plasma. Type 1 interferons have previously been proposed as an accurate diagnostic screening test for pediatric viral infection.^36^ Another diagnostic challenge in vLRTI is identifying bacterial coinfection, as standard respiratory bacterial cultures do not distinguish between coinfection and colonization and are often negative in the context of prior antibiotic administration. Our subanalysis of bacterial coinfection highlighted two TA proteins, TSG-6 (a tumor necrosis factor-inducible protein) and CRP (an inflammatory protein with modest specificity for bacterial LRTI in blood),^37^ that may be useful respiratory biomarkers of secondary bacterial infection.

Our study has several strengths including our multi-center enrollment, clinical sampling at early time points, evaluation of protein expression in multiple compartments, and integration of respiratory mNGS for comprehensive pathogen evaluation. It also has several important limitations including small sample size which may have limited our ability to detect more subtle but clinically important differences in protein expression. Additionally, the version of the SomaScan^®^ assay utilized does not encompass the entire human proteome, and we likely missed some important differentially expressed proteins and pathways. From the diagnostic biomarker standpoint, our findings are more preliminary in nature, and warrant further optimization and validation in larger cohorts with all relevant classes of infection (bacterial infection, viral infection, coinfection, and non-infectious controls) and a wider range of severity represented to rigorously understand performance. Finally, we recognize that there is no gold standard for LRTI diagnosis in children, and our reliance on the best practical methodology of combining retrospective clinical adjudication and microbiology results may have resulted in classification errors.

Taken together, we present a comprehensive proteomic characterization of severe pediatric vLRTI, highlighting pathophysiologic insights in both viral infection and bacterial-viral coinfection and deepening our understanding of compartmentalization of the human host response to LRTI. Validation of the present findings in larger external cohorts are needed with more in-depth analysis to determine whether new therapeutic targets can be identified and whether proteomic biomarkers may augment current standard-of-care pathogen-based diagnostic testing. Looking forward, multi-omic approaches combining proteomics and transcriptomics as well as integration with microbiology hold promise for advancing understanding of the heterogeneity of pediatric LRTI, modernizing diagnostics, and personalizing treatment.

## DATA AVAILABILITY

Proteomic data, subject metadata, and code for reproducing the results of this study can be found at: https://github.com/infectiousdisease-langelier-lab/pedsLRTIproteomics.

## REFERENCES

1 Hemming VG. Viral respiratory diseases in children: Classification, etiology, epidemiology, and risk factors. The Journal of Pediatrics 1994; 124: S13.

2 Liu L, Oza S, Hogan D, et al. Global, regional, and national causes of under-5 mortality in 2000–15: an updated systematic analysis with implications for the Sustainable Development Goals. Lancet 2016; 388: 3027–35.

3 Rouse BT, Sehrawat S. Immunity and immunopathology to viruses: what decides the outcome? Nat Rev Immunol 2010; 10: 514–26.

4 Zaas AK, Garner BH, Tsalik EL, Burke T, Woods CW, Ginsburg GS. The current epidemiology and clinical decisions surrounding acute respiratory infections. Trends Mol Med 2014; 20: 579–88.

5 Troy NM, Bosco A. Respiratory viral infections and host responses; insights from genomics. Respiratory Research 2016; 17: 156.

6 Wildman E, Mickiewicz B, Vogel HJ, Thompson GC. Metabolomics in pediatric lower respiratory tract infections and sepsis: a literature review. Pediatr Res 2023; 93: 492–502.

7 Casini F, Valentino MS, Lorenzo MG, et al. Use of transcriptomics for diagnosis of infections and sepsis in children: A narrative review. Acta Paediatrica 2024; 113: 670–6.

8 Mick E, Tsitsiklis A, Kamm J, et al. Integrated host/microbe metagenomics enables accurate lower respiratory tract infection diagnosis in critically ill children. J Clin Invest 2023; 133. DOI:10.1172/JCI165904.

9 Mourani PM, Sontag MK, Williamson KM, et al. Temporal airway microbiome changes related to ventilator-associated pneumonia in children. European Respiratory Journal 2021; 57. DOI:10.1183/13993003.01829-2020.

10 Dapat C, Kumaki S, Sakurai H, et al. Gene signature of children with severe respiratory syncytial virus infection. Pediatr Res 2021; 89: 1664–72.

11 Pereira-Fantini PM, Tingay DG. The proteomics of lung injury in childhood: challenges and opportunities. Clin Proteomics 2016; 13: 5.

12 Cheng J, Ji D, Yin Y, et al. Proteomic profiling of urinary small extracellular vesicles in children with pneumonia: a pilot study. Pediatr Res 2023; 94: 161–71.

13 Tsai M-H, Lin T-Y, Hsieh S-Y, Chiu C-Y, Chiu C-H, Huang Y-C. Comparative proteomic studies of plasma from children with pneumococcal pneumonia. Scandinavian Journal of Infectious Diseases 2009; 41: 416–24.

14 Yin G-Q, Zeng H-X, Li Z-L, et al. Differential proteomic analysis of children infected with respiratory syncytial virus. Braz J Med Biol Res 2021; 54: e9850.

15 Gold L, Ayers D, Bertino J, et al. Aptamer-based multiplexed proteomic technology for biomarker discovery. PLoS One 2010; 5: e15004.

16 Kim CH, Tworoger SS, Stampfer MJ, et al. Stability and reproducibility of proteomic profiles measured with an aptamer-based platform. Sci Rep 2018; 8: 8382.

17 Candia J, Cheung F, Kotliarov Y, et al. Assessment of Variability in the SOMAscan Assay. Sci Rep 2017; 7: 14248.

18 Ritchie ME, Phipson B, Wu D, et al. limma powers differential expression analyses for RNA-sequencing and microarray studies. Nucleic Acids Res 2015; 43: e47.

19 Milacic M, Beavers D, Conley P, et al. The Reactome Pathway Knowledgebase 2024. Nucleic Acids Research 2024; 52: D672–8.

20 Wang J, Vasaikar S, Shi Z, Greer M, Zhang B. WebGestalt 2017: a more comprehensive, powerful, flexible and interactive gene set enrichment analysis toolkit. Nucleic Acids Research 2017; 45: W130–7.

21 Tay JK, Narasimhan B, Hastie T. Elastic Net Regularization Paths for All Generalized Linear Models. Journal of Statistical Software 2023; 106: 1–31.

22 Mesev EV, LeDesma RA, Ploss A. Decoding type I and III interferon signalling during viral infection. Nat Microbiol 2019; 4: 914–24.

23 de Jong LC, Crnko S, ten Broeke T, Bovenschen N. Noncytotoxic functions of killer cell granzymes in viral infections. PLoS Pathog 2021; 17: e1009818.

24 Jin R, Hao J, Yi Y, Sauter E, Li B. Regulation of macrophage functions by FABP-mediated inflammatory and metabolic pathways. Biochim Biophys Acta Mol Cell Biol Lipids 2021; 1866: 158964.

25 Storch J, Thumser AE. Tissue-specific Functions in the Fatty Acid-binding Protein Family. J Biol Chem 2010; 285: 32679–83.

26 Connolly E, Hussell T. The Impact of Type 1 Interferons on Alveolar Macrophage Tolerance and Implications for Host Susceptibility to Secondary Bacterial Pneumonia. Front Immunol 2020; 11: 495.

27 Mehta D, Petes C, Gee K, Basta S. The Role of Virus Infection in Deregulating the Cytokine Response to Secondary Bacterial Infection. J Interferon Cytokine Res 2015; 35: 925–34.

28 Bai M, Grieshaber-Bouyer R, Wang J, et al. CD177 modulates human neutrophil migration through activation-mediated integrin and chemoreceptor regulation. Blood 2017; 130: 2092–100.

29 de Kretser DM, Bensley JG, Pettilä V, et al. Serum activin A and B levels predict outcome in patients with acute respiratory failure: a prospective cohort study. Crit Care 2013; 17: 1–13.

30 Devanathan V, Hagedorn I, Köhler D, et al. Platelet Gi protein Gαi2 is an essential mediator of thrombo-inflammatory organ damage in mice. Proc Natl Acad Sci U S A 2015; 112: 6491–6.

31 Dunning J, Blankley S, Hoang LT, et al. Progression of whole-blood transcriptional signatures from interferon-induced to neutrophil-associated patterns in severe influenza. Nat Immunol 2018; 19: 625–35.

32 Novak T, Crawford JC, Hahn G, et al. Transcriptomic profiles of multiple organ dysfunction syndrome phenotypes in pediatric critical influenza. Front Immunol 2023; 14: 1220028.

33 Majewski S, Zhou X, Miłkowska-Dymanowska J, Białas AJ, Piotrowski WJ, Malinovschi A. Proteomic profiling of peripheral blood and bronchoalveolar lavage fluid in interstitial lung diseases: an explorative study. ERJ Open Research 2021; 7. DOI:10.1183/23120541.00489-2020.

34 García-Arroyo L, Prim N, Martí N, Roig MC, Navarro F, Rabella N. Benefits and drawbacks of molecular techniques for diagnosis of viral respiratory infections. Experience with two multiplex PCR assays. J Med Virol 2016; 88: 45–50.

35 Jansen RR, Wieringa J, Koekkoek SM, et al. Frequent Detection of Respiratory Viruses without Symptoms: Toward Defining Clinically Relevant Cutoff Values ▿. J Clin Microbiol 2011; 49: 2631–6.

36 Trouillet-Assant S, Viel S, Ouziel A, et al. Type I Interferon in Children with Viral or Bacterial Infections. Clinical Chemistry 2020; 66: 802–8.

37 van der Meer V, Neven AK, van den Broek PJ, Assendelft WJJ. Diagnostic value of C reactive protein in infections of the lower respiratory tract: systematic review. BMJ 2005; 331: 26.

